# Near Perfect Neural Critic from Motor Cortical Activity Toward an Autonomously Updating Brain Machine Interface

**DOI:** 10.1101/250316

**Authors:** Junmo An, Taruna Yadav, Mohammad Badri Ahmadi, Venkata S Aditya Tarigoppula, Joseph Thachil Francis

## Abstract

We are developing an autonomously updating brain machine interface (BMI) utilizing reinforcement learning principles. One aspect of this system is a neural critic that determines reward expectations from neural activity. This critic is then used to update a BMI decoder towards an improved performance from the user’s perspective. Here we demonstrate the ability of a neural critic to classify trial reward value given activity from the primary motor cortex (M1), using neural features from single/multi units (SU/MU), and local field potentials (LFPs) with prediction accuracies *up to 97% correct*. A nonhuman primate subject conducted a cued center out reaching task, either manually, or observationally. The cue indicated the reward value of a trial. Features such as power spectral density (PSD) of the LFPs and spike-field coherence (SFC) between SU/MU and corresponding LFPs were calculated and used as inputs to several classifiers. We conclude that hybrid features of PSD and SFC show higher classification performance than PSD or SFC alone (accuracy was 92% for manual tasks, and 97% for observational). In the future, we will employ these hybrid features towards our autonomously updating BMI.

## I. Introduction

Over the past several years, brain machine interface (BMI) technology has grown significantly to help research participants with motor disabilities control computer cursors [1] or robotic prosthetic limbs [2]. Traditionally, BMI decoders have been trained using supervised learning techniques to translate movement-related neural information from the motor cortex. However, supervised learning based BMIs require an explicit training set of both neural and kinematic data to learn the BMI output commands [3], and thus can’t easily be updated in real world use situations.

To address this challenge, we are currently developing a reinforcement learning (RL) based BMI that can update autonomously using an evaluative feedback signal provided by a neurally derived critic [4-9]. Such an RL-BMI’s performance is directly dependent on the fidelity of the neural critic, and therefore accurate classification of neural activity by the critic is helpful. To advance the neural critic accuracy, we used frequency domain features including power spectral density (PSD) from an ensemble’s averaged local field potential (LFPs), and spike-field coherence (SFC) to classify the neural activity in the primary motor cortex (M1) from center-out reaching tasks differing in reward expectation on a trial-by-trial basis. PSD of LFPs have been used extensively for neural data analysis, and SFC has been used for cross-scale interaction analysis between the microscale, spikes, and macroscale, LFPs [10-12]. We tested classification performance on different conventional classifiers and feature sets, and found that hybrid features (PSD and SFC) achieved a higher classification accuracy than individual PSD or SFC features. The use of real-time computable hybrid features resulted in a near perfect classification accuracy and should be advantageous in the autonomous updating of the online BMI control.

## II. Methods

### A. Psychophysical Tasks and Neural Data Recording

One macaque monkey was implanted in the left M1. We used 96-channel platinum microelectrode arrays (10 x 10 Utah array with 1.5 mm deep electrodes separated by 400 μm, Blackrock Microsystems) to simultaneously record single-unit (spikes) and LFPs. All procedures were approved by SUNY Downstate Medical Center’s (IACUC) and complied with the National Institutes of Health Guide for the Care and Use of Laboratory Animals.

The monkey was trained on two tasks, manual and observational. For the manual task, the monkey performed one target center-out reaching movements with its right arm resting comfortably inside a two-joint exoskeletal robot (KINARM system, BKIN Technologies), Figure 1(a). During the observational task, the monkey passively observed the movement of a feedback cursor from the center to the peripheral target on the display screen while the KINARM was locked in place, Figure 1(b). Each trial consisted of four events, center hold, cue onset, reach to target and target hold. For the manual task, the monkey initiated a trial by fixating and holding the center target until the color cued peripheral reach target indicating the rewarding or non-rewarding outcomes was presented. Next, the monkey waited for 300 ms until the color-cued center target disappeared (representing an implicit GO cue). To complete the trial successfully, the monkey had to hold on the peripheral target position for 325 ms following a successful reach to the target. Every successful reach for a cued rewarding trial ended with a juice reward while no reward was given for successful cued non-rewarding trials. To encourage reaching movements in all trials, an unsuccessful cued non-rewarding trial was always followed by another non-rewarding trial. Similar to the manual task, visual color cues were presented during the observational task. The order of rewarding and non-rewarding trials was randomly chosen in the manual tasks whereas the reward and non-rewarding trials were presented in a predictable sequence, reward followed by non-rewarding trial repeated, during the observational tasks [6].

**Figure 1.**
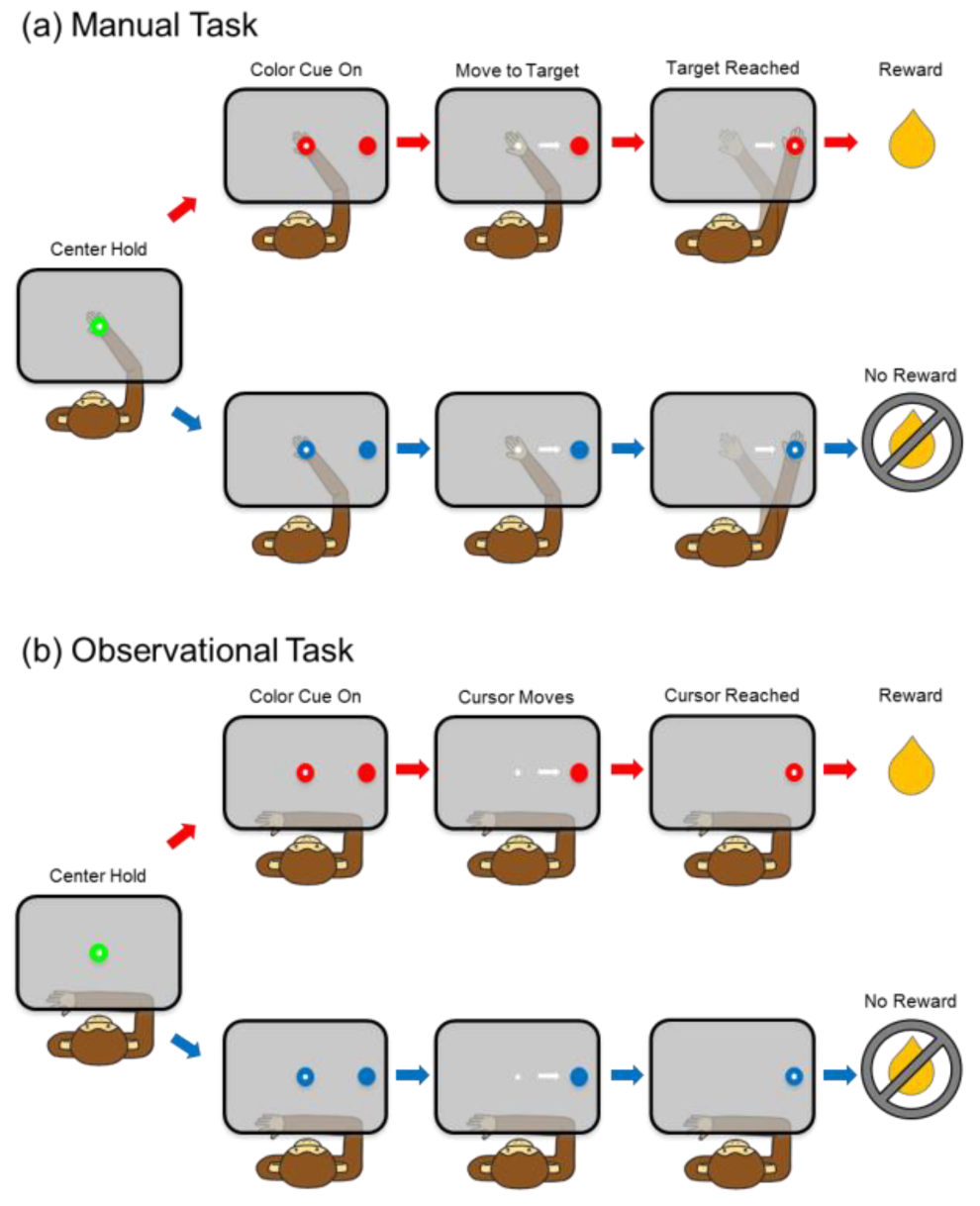
Schematic of the (a) manual task and (b) observational task.

Multichannel acquisition processor systems (Plexon Inc.) were used to simultaneously record spiking activity and LFPs from contralateral M1 at a sampling rate of 40 KHz and 2000 Hz respectively. The composite signal (consisting of spikes and LFPs) were pre-amplified and band-pass filtered from 0.5 to 300 Hz for LFPs and 170 Hz to 8 kHz for single-unit activity. We recorded from 32 LFP channels and 174 units (80 units for manual task and 94 units for observational task) in M1 while the monkey performed/observed the center-out reaching tasks.

### B. Classification Procedure

Figure 2 shows the flowchart for the proposed classification process using PSD, SFC and hybrid features: (1) recording of spike trains and corresponding LFPs, (2) calculation of PSD and SFC, respectively, (3) applying normalization, (4) reducing dimensionality of PSD, SFC, and hybrid features, (5) classification and cross-validation, (6) comparing classification accuracies. The preprocessing for feature extraction was performed with MATLAB, and the classification and validation were implemented in Python with TensorFlow and Scikit-Learn machine learning libraries.

**Figure 2.**
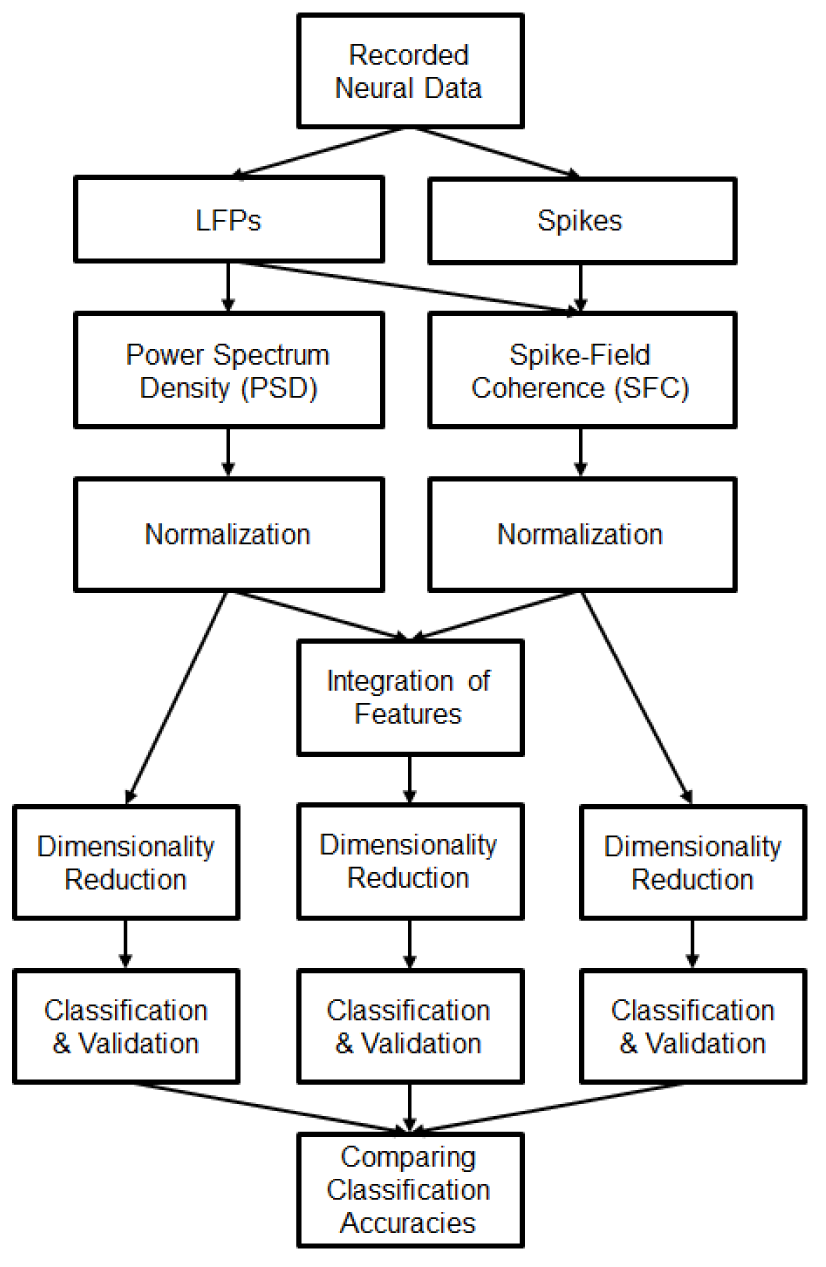
Flowchart of the proposed classification procedure using hybrid features.

### C. Feature Selection and Extraction

PSD and SFC were measured and used as hybrid features to discriminate between rewarding and non-rewarding trials. For both PSD and SFC measures, we used a post-cue-onset period of 800 ms for the manual task and 2 seconds for the observational task. Before computing PSD of the LFPs, 60 Hz line noise and harmonics were eliminated using a second-order notch filter. For analyses, we only selected LFP channels with signal-to-noise ratios (SNR) ≥4. After normalizing each LFP channel using min-max normalization method, the SNR was calculated for each channel as the ratio of peak-to-peak amplitude of averaged LFP and twice the standard deviation of the residual signal (given by the difference of i^th^ channel’s LFP and the averaged LFP) as shown in (1) [13]

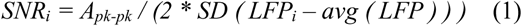

We then averaged the selected LFP channels which were later used for PSD and SFC computations. The PSD was computed from LFP traces using the Welch periodogram method [14] with 75% overlapping windows. Normalization was applied to each power spectrum by dividing power at each frequency by the sum of all power from 0.5 to 100 Hz [15].

SFC has been used to measure phase relationships between spikes and LFPs. SFC was computed using the coherence equation shown in (2) [10-12]

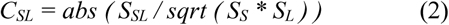

Equation 2 shows that the *CSL* can be calculated as the ratio of *S_SL_* (cross spectrum between spikes and LFPs) and the square root of the product of spectra of spikes (*S_S_*) and LFPs (*S_L_*). SFC (*C_SL_*) returns a value that ranges from 0 to 1. SFC value of 1 indicates perfect phase dependency between spikes and LFPs, and a value 0 means no phase dependency between them.

The extracted features such as PSD and SFC were stored in a feature matrix. In this matrix, each row corresponds to each trial during both rewarding and non-rewarding trials, whereas columns correspond to features. To configure hybrid features, all attributes of PSD features were shifted and rescaled to the same ranges of SFC features (ranging from 0 to 1). PSD and SFC features were then concatenated.

For classification, PSD, SFC, and integrated features were reduced to two dimensions by transforming the data using one of two dimensionality reduction techniques. We either integrated linear and nonlinear dimensionality reduction (also known as manifold learning), or used manifold learning alone. Principal Component Analysis (PCA) was used to reduce the feature’s dimensionality with an explained variance ratio of 95% first and then Multidimensional Scaling (MDS) was applied [16, 17], or we used a t-Distributed Stochastic Neighbor Embedding (t-SNE) [18] method alone, as these two methods consistently outperformed others we tested. We chose the best results from these two methods for Tables 1 and 2, as indicated.

**Table 1.** CLASSIFICATION ACCURACY FOR MANUAL TASKS (AVERAGE ± STANDARD DEVIATION). (I): PCA+MDS, (II): T-SNE.

| Classifiers | Features (%) | Accuracy (%) | Sensitivity (%) | Specificity (%) |
| --- | --- | --- | --- | --- |
| **MLP** | PSD (I) | 76 ± 10 | 79 ± 10 | 78 ± 8 |
| | SFC (I) | 83 ± 16 | 88 ± 14 | 83 ± 12 |
|  | **Hybrid (I)** | **92 ± 10** | **92 ± 8** | **92 ± 9** |
| kNN | PSD (II) | 75 ± 13 | 76 ± 15 | 79 ± 14 |
| | SFC (II) | 73 ± 15 | 77 ± 17 | 77 ± 18 |
| | Hybrid (II) | 78 ± 12 | 75 ± 9 | 79 ± 16 |
| RBF SVM | PSD (II) | 75 ± 9 | 77 ± 12 | 80 ± 5 |
| | SFC (II) | 72 ± 16 | 77 ± 13 | 76 ± 12 |
| | Hybrid (II) | 78 ± 10 | 81 ± 9 | 79 ± 14 |
| Naive Bayes | PSD (I) | 81 ± 5 | 79 ± 8 | 83 ± 11 |
| | SFC (I) | 82 ± 16 | 80 ± 16 | 84 ± 13 |
| | Hybrid (I) | 85 ± 11 | 82 ± 8 | 84 ± 9 |
| Gaussian Process | PSD (II) | 80 ± 8 | 78 ± 5 | 79 ± 4 |
| | SFC (II) | 80 ± 15 | 79 ± 16 | 80 ± 11 |
| | Hybrid (II) | 86 ± 8 | 89 ± 16 | 87 ± 10 |

**Table 2.**
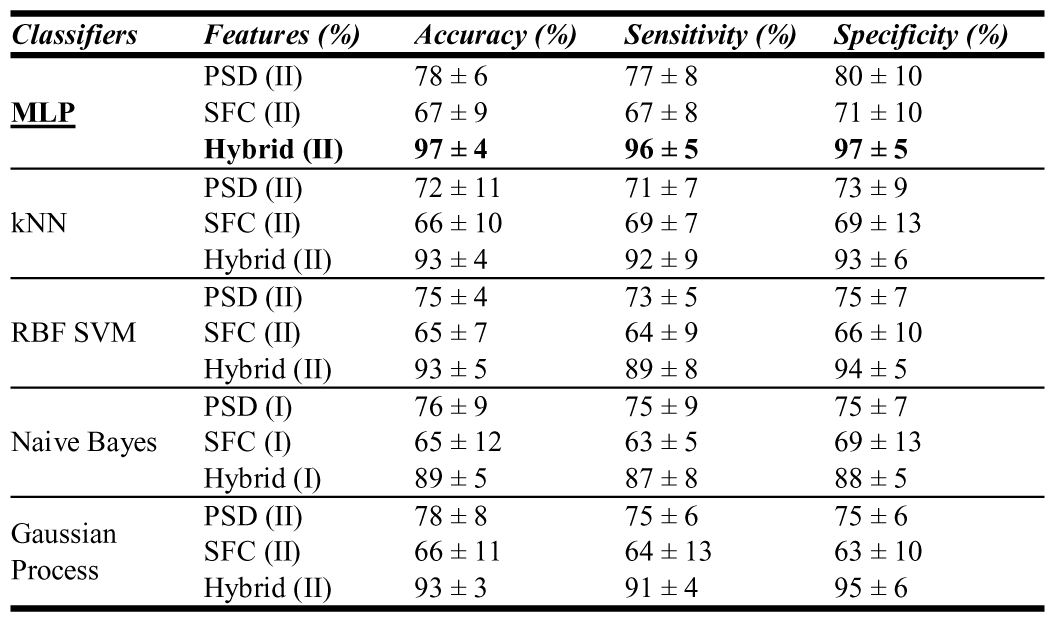
CLASSIFICATION ACCURACY FOR OBSERVATIONAL TASKS (AVERAGE ± STANDARD DEVIATION). (I): PCA+MDS, (II): T-SNE.

| Classifiers | Features (%) | Accuracy (%) | Sensitivity (%) | Specificity (%) |
| --- | --- | --- | --- | --- |
| **MLP** | PSD (II) | 78 ± 6 | 77 ± 8 | 80 ± 10 |
| | SFC (II) | 67 ± 9 | 67 ± 8 | 71 ± 10 |
|  | **Hybrid (II)** | **97 ± 4** | **96 ± 5** | **97 ± 5** |
| kNN | PSD (II) | 72 ± 11 | 71 ± 7 | 73 ± 9 |
| | SFC (II) | 66 ± 10 | 69 ± 7 | 69 ± 13 |
| | Hybrid (II) | 93 ± 4 | 92 ± 9 | 93 ± 6 |
| RBF SVM | PSD (II) | 75 ± 4 | 73 ± 5 | 75 ± 7 |
| | SFC (II) | 65 ± 7 | 64 ± 9 | 66 ± 10 |
| | Hybrid (II) | 93 ± 5 | 89 ± 8 | 94 ± 5 |
| Naive Bayes | PSD (I) | 76 ± 9 | 75 ± 9 | 75 ± 7 |
| | SFC (I) | 65 ± 12 | 63 ± 5 | 69 ± 13 |
| | Hybrid (I) | 89 ± 5 | 87 ± 8 | 88 ± 5 |
| Gaussian Process | PSD (II) | 78 ± 8 | 75 ± 6 | 75 ± 6 |
| | SFC (II) | 66 ± 11 | 64 ± 13 | 63 ± 10 |
| | Hybrid (II) | 93 ± 3 | 91 ± 4 | 95 ± 6 |

### D. Classification

The extracted feature vectors that are obtained from PSD, SFC methods and their hybrid were given as input to widely used classifiers, such as Multi-Layer Perceptron Network (MLP), k-Nearest Neighbors (kNN), Radial Basis Function kernel Support Vector Machines (RBF SVM), Gaussian Naive Bayes and Gaussian Process. The MLP classifier was composed of three layers of neurons, an input layer, a hidden layer and an output layer. The hidden layer was composed of 100 ReLU (rectified linear) neurons. Stochastic gradient descent optimizer for MLP was used to minimize the loss function. The kNN classifier was trained for 3 neighbors on the training set. The parameters of the RBF SVM were assigned as gamma = 2 and C = 1. Gaussian Naive Bayes and Gaussian Process were applied with default parameter settings as suggested by Scikit-Learn.

### E. Validation

In order to validate the proposed classification, a ten-fold cross-validation was performed for splitting datasets into ten consecutive folds. Each fold was used once as a validation set, while the rest of the folds were used for training. In addition, the performance of classification, such as accuracies, sensitivities, and specificities were computed, as given in the Equations (3), (4) and (5).

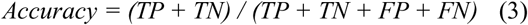

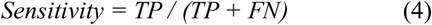

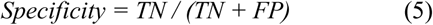

Here TP, TN, FP and FN denote the number of true positives, true negatives, false positives, and false negatives, respectively.

## III. Results

In order to classify reward expectation from neural activity in M1 into rewarding and non-rewarding trials in the post-cue period, neural features such as PSD, SFC and their hybrid were computed, and applied as input data to different conventional classifiers such as MLP, kNN, RBF SVM,

Gaussian Naive Bayes and Gaussian Process. The classification performance was then calculated by applying accuracy, sensitivity and specificity using ten-fold cross-validation. Finally, we investigated the comparison of classification performance calculated across five different classifiers and each feature set.

Table 1 and 2 represent the classification comparison for manual and observational tasks, respectively. For manual tasks, PCA+MDS method was used for MLP and Naive Bayes classifiers whereas t-SNE method was used for the other three classifiers. For observational tasks, we used PCA+MDS for Naive Bayes classifier whereas t-SNE was used for all other classifiers. The results, as shown in Table 1, indicate that hybrid of PSD and SFC features for the manual task produces higher classification performance than individual PSD or SFC features. The MLP using hybrid features yielded higher classification performance (accuracy, sensitivity and specificity are all 92%) as compared to other classifiers. Similar to the classification results for the manual task, the MLP using hybrid features for the observational task also achieved the highest classification performance (accuracy = 97%, sensitivity = 96%, and specificity = 97%) as shown in Table 2.

We also compared the classification accuracy of our classifiers with the PSD, SFC or their hybrid (reduced dimension) features as inputs based on receiver operating characteristic (ROC) curves as well as the area under the ROC curve (AUC). Figure 3 displays the ROC curves with AUC values for the MLP using PSD, SFC, and hybrid for manual (a) and observational (b) tasks (ROC curves for other classifiers not shown). As shown in Fig. 3, the AUCs of the ROC curve of MLP using hybrid features are larger than using individual PSD or SFC features for both manual and observational tasks.

**Figure 3.**
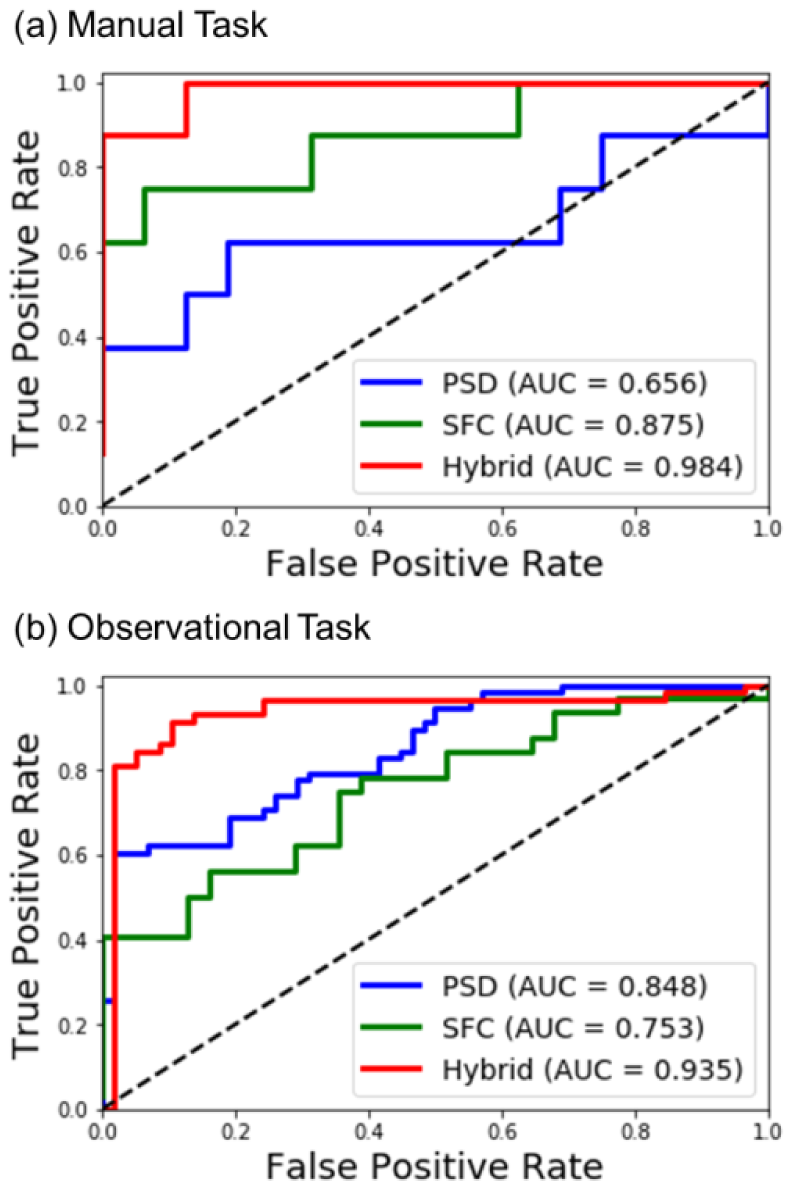
ROC curves with AUC values of MLP methods for testing set using PSD (blue), SFC (green), and Hybrid (red) for (a) manual and (b) observational task.

## IV. Discussion and Conclusion

The results of this study indicate that reward expectation from M1 neural activity can discriminate between rewarding and non-rewarding trials during both reaching movements (manual tasks) and passive observations (observational tasks) [6]. We showed, using common classifiers, that integration of neural features such as PSD and SFC provide higher classification accuracy than either one individually. Moreover, a relatively simple MLP classifier yielded the highest classification performance as compared to other classifiers. We also found that choosing one dimensionality reduction method over the other was a tradeoff between computation time and classification performance. Average classification accuracy using simple PCA method was slightly worse (3 to 5% less) than using PCA+MDS or t-SNE. However, the computation time of PCA+MDS or t-SNE was greater than that of the PCA method. In future investigations, it might be possible to use different types of deep neural network architectures that can yield more robust, reliable classification performance to decode movement-related information from the neural signals, including movement initiation and therefore autonomous windowing for the above feature extraction.

In conclusion, we tested and validated the classification to decode reward-related neural signals in M1 during both manual and observational tasks. This classification technique using hybrid PSD and SFC features should be useable as a neural critic to quickly adapt an RL-BMI to better decode the user's intended movements in all environments.

## References

[1] J. R. Wolpaw and D. J. McFarland, “Control of a two-dimensional movement signal by a noninvasive brain-computer interface in humans”, Proceedings of the National Academy of Sciences of the United States of America, vol. 101, pp. 17849–17854, 2004.

[2] L. R. Hochberg, D. Bacher, B. Jarosiewicz, N. Y. Masse, J. D. Simeral, J. Vogel, et al., “Reach and grasp by people with tetraplegia using a neurally controlled robotic arm”, Nature, vol. 485, pp. 372–375, 2012.

[3] V. Gilja, P. Nuyujukian, C. A. Chestek, J. P. Cunningham, M. Y. Byron, J. M. Fan, et al., “A high-performance neural prosthesis enabled by control algorithm design”, Nature neuroscience, vol. 15, pp. 1752–1757, 2012.

[4] N. W. Prins, J. C. Sanchez, and A. Prasad, “A confidence metric for using neurobiological feedback in actor-critic reinforcement learning based brain-machine interfaces”, Frontiers in neuroscience, vol. 8, 2014.

[5] J. DiGiovanna, B. Mahmoudi, J. Fortes, J. C. Principe, and J. C. Sanchez, “Coadaptive brain machine interface via reinforcement learning”, IEEE transactions on biomedical engineering, vol. 56, pp. 54–64, 2009.

[6] B. T. Marsh, V. S. A. Tarigoppula, C. Chen, and J. T. Francis, “Toward an autonomous brain machine interface: integrating sensorimotor reward modulation and reinforcement learning”, Journal of Neuroscience, vol. 35, pp. 7374–7387, 2015.

[7] J. C. Sanchez, B. Mahmoudi, J. DiGiovanna, and J. C. Principe, “Exploiting co-adaptation for the design of symbiotic neuroprosthetic assistants”, Neural Networks, vol. 22, pp. 305–315, 2009.

[8] A. Tarigoppula, N. Rotella, and J. T. Francis, “Properties of a temporal difference reinforcement learning brain machine interface driven by a simulated motor cortex”, in Engineering in Medicine and Biology Society (EMBC), 2012 Annual International Conference of the IEEE, 2012, pp. 3284–3287.

[9] D. B. McNiel, J. S. Choi, J. P. Hessburg, and J. T. Francis, “Reward value is encoded in primary somatosensory cortex and can be decoded from neural activity during performance of a psychophysical task”, in Engineering in Medicine and Biology Society (EMBC), 2016 IEEE 38th Annual International Conference of the, 2016, pp. 3064–3067.

[10] P. Fries, J. H. Reynolds, A. E. Rorie, and R. Desimone, “Modulation of oscillatory neuronal synchronization by selective visual attention”, Science, vol. 291, pp. 1560–1563, 2001.

[11] P. Fries, P. R. Roelfsema, A. K. Engel, P. König, and W. Singer, “Synchronization of oscillatory responses in visual cortex correlates with perception in interocular rivalry”, Proceedings of the National Academy of Sciences, vol. 94, pp. 12699–12704, 1997.

[12] M. Jarvis and P. Mitra, “Sampling properties of the spectrum and coherency of sequences of action potentials”, Neural Computation, vol. 13, pp. 717–749, 2001.

[13] A. C. Koralek, R. M. Costa, and J. M. Carmena, “Temporally precise cell-specific coherence develops in corticostriatal networks during learning”, Neuron, vol. 79, pp. 865–872 2013.

[14] P. Welch, “The use of fast Fourier transform for the estimation of power spectra: a method based on time averaging over short, modified periodograms”, IEEE Transactions on audio and electroacoustics, vol. 15, pp. 70–73, 1967.

[15] L. L. Colgin, T. Denninger, M. Fyhn, T. Hafting, T. Bonnevie, O. Jensen, et al., “Frequency of gamma oscillations routes flow of information in the hippocampus”, Nature, vol. 462, pp. 353–357, 2009.

[16] A. Géron, Hands-on machine learning with Scikit-Learn and TensorFlow: concepts, tools, and techniques to build intelligent systems, ed: O’Reilly Media, Inc, 2017.

[17] J. B. Kruskal, “Multidimensional scaling by optimizing goodness of fit to a nonmetric hypothesis”, Psychometrika, vol. 29, pp. 1–27, 1964.

[18] L. v. d. Maaten and G. Hinton, “Visualizing data using t-SNE”, Journal of Machine Learning Research, vol. 9, pp. 2579–2605, 2008.

